# Language barriers in global bird conservation

**DOI:** 10.1101/2021.05.24.445290

**Authors:** Pablo Jose Negret, Scott C. Atkinson, Bradley K. Woodworth, Marina Corella Tor, James R. Allan, Richard A. Fuller, Tatsuya Amano

**Affiliations:** The University of Queensland, School of Earth and Environmental Sciences, Qld 4072, Australia; The University of Queensland, Centre for Biodiversity and Conservation Science, Qld 4072, Australia; United Nations Development Programme, New York, U.S.A; The University of Queensland, School of Biological Sciences, Qld 4072, Australia; Institute for Biodiversity and Ecosystem Dynamics, University of Amsterdam, Amsterdam, The Netherlands

## Abstract

Multiple languages being spoken within a species’ distribution can impede communication among conservation stakeholders, the compilation of scientific information, and the development of effective conservation actions. Here, we investigate the number of official languages spoken within the distributions of 10,863 bird species to identify which ones might be particularly affected by consequences of language barriers. We show that 1587 species have 10 languages or more spoken within their distributions. Threatened, migratory and wide-ranging species have especially many languages spoken within their distribution. Particularly high numbers of species with many languages within their distribution are found in Eastern Europe, Russia and central and western Asia. Global conservation efforts would benefit from implementing guidelines to overcome language barriers, especially in regions with high species and language diversity.

## Introduction

Earth’s biodiversity is under threat. Human population growth and associated activities are causing the loss of natural ecosystems and species habitats at an unprecedented rate (*1, 2*), with at least one million species currently threatened with extinction (*3*). This accelerated loss of biodiversity and the fact that many species and threats extend beyond country borders has stimulated the generation of guidelines for effective transboundary collaboration on international agreements, such as the Convention on Biological Diversity and the Convention on Trade in Endangered Species of Wild Fauna and Flora (*4, 5*). However, existing guidelines for transboundary collaboration rarely consider differences in cultural backgrounds among countries, which can create both challenges and opportunities in conservation (*4, 6, 7*).

An aspect of culture that has fundamental consequences for conservation is the variety of languages that people speak. Language differences across the distribution of a species can generate a number of challenges for conservation (summarized with examples in Table 1). First, multiple languages being spoken within the distribution of a species can create a barrier to the effective collection and compilation of scientific information relevant to conservation, which is often scattered across languages (*6*). For example, comprehensive ecological knowledge of understudied seasonal migratory birds in Brazil could only be achieved by combining information from Brazilian citizen science platforms available only in Portuguese with information from global, English-language, platforms (7). Second, language differences within the distribution of species can also impede effective agreements between stakeholders in conservation decisions. For example, differences in the use of vocabulary even within the same language influenced the perception of the public on the importance of hedgehog eradication as a conservation measure in Scotland (*9*). Such an effect could be magnified further when stakeholders speak different languages. Third, language differences can affect the generation and quality of collaborative conservation projects. For example, overcoming language barriers was recognized as a fundamental step for the generation of effective conservation measures for threatened bird species in the Julian alps, the Bavarian-Bohemian Forest (*10*) and the Mediterranean sea (*11*).

**Table 1.**
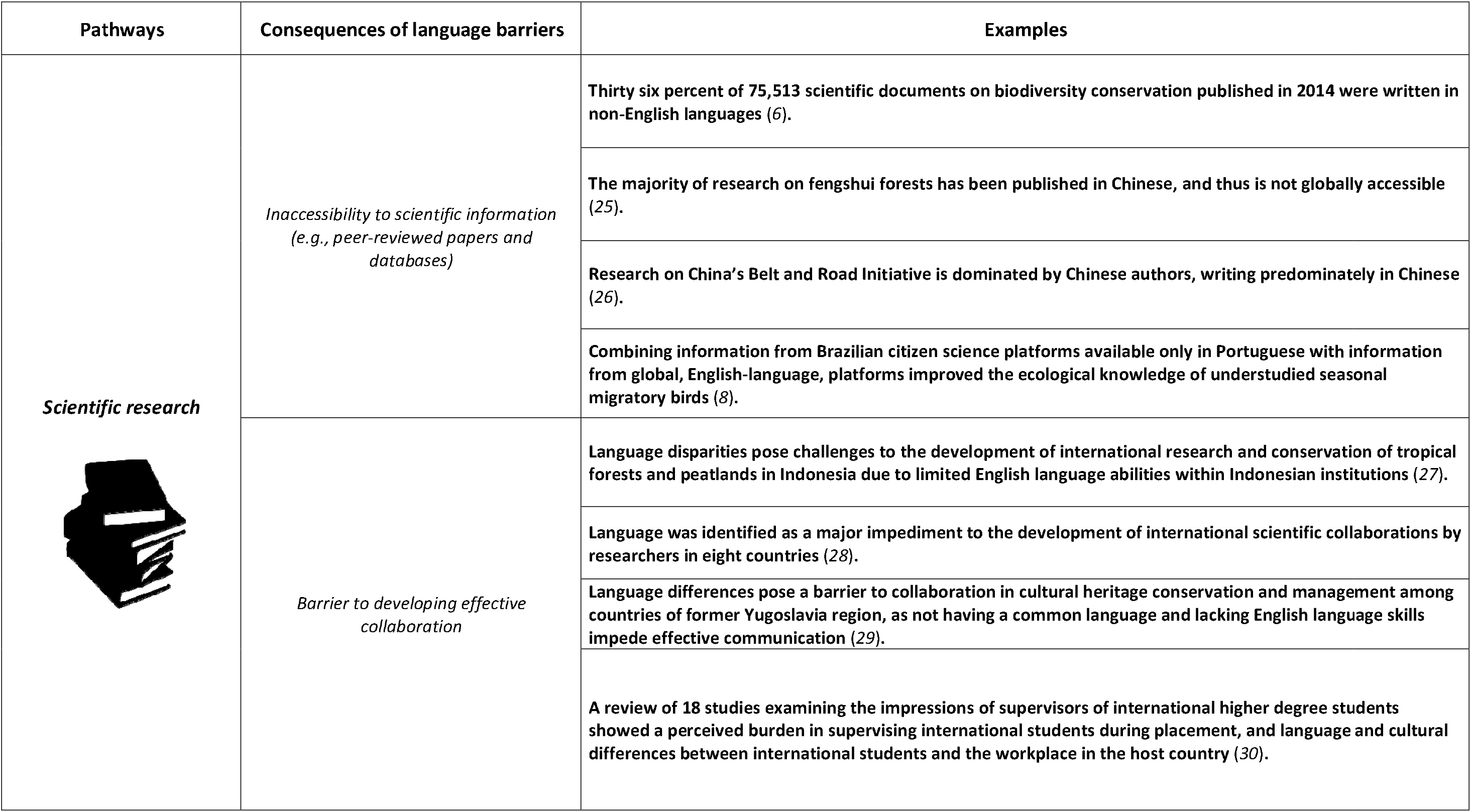
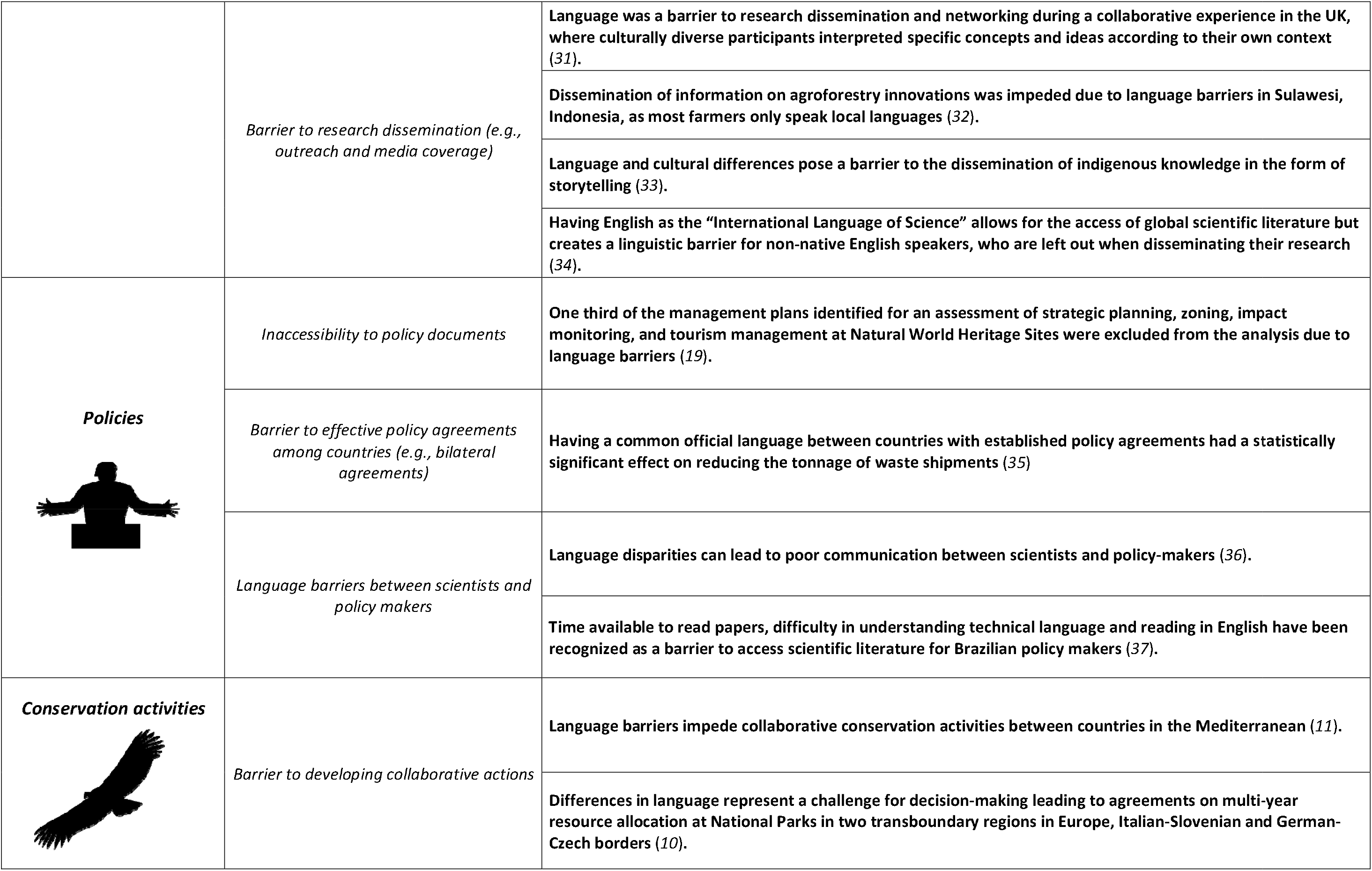
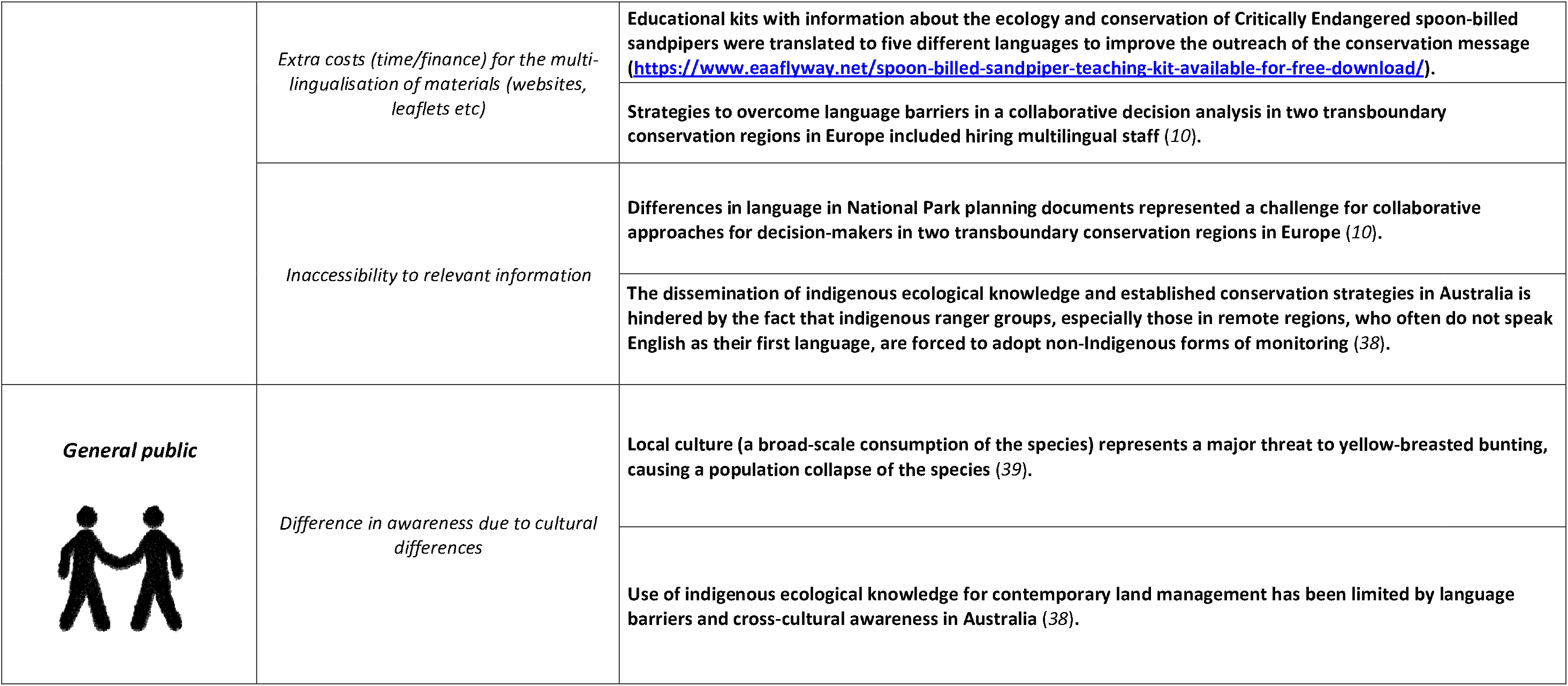
Potential challenges to conservation outcomes caused by language barriers

| Pathways | Consequences of language barriers | Examples |
| --- | --- | --- |
| Scientific research | *Inaccessibility to scientific information (e.g., peer-reviewed papers and databases)* | Thirty six percent of 75,513 scientific documents on biodiversity conservation published in 2014 were written in non-English languages (6). |
|  |  | The majority of research on fengshui forests has been published in Chinese, and thus is not globally accessible (25). |
|  |  | Research on China’s Belt and Road Initiative is dominated by Chinese authors, writing predominately in Chinese (26). |
|  |  | Combining information from Brazilian citizen science platforms available only in Portuguese with information from global, English-language, platforms improved the ecological knowledge of understudied seasonal migratory birds (8). |
|  | *Barrier to developing effective collaboration* | Language disparities pose challenges to the development of international research and conservation of tropical forests and peatlands in Indonesia due to limited English language abilities within Indonesian institutions (27). |
|  |  | Language was identified as a major impediment to the development of international scientific collaborations by researchers in eight countries (28). |
|  |  | Language differences pose a barrier to collaboration in cultural heritage conservation and management among countries of former Yugoslavia region, as not having a common language and lacking English language skills impede effective communication (29). |
|  |  | A review of 18 studies examining the impressions of supervisors of international higher degree students showed a perceived burden in supervising international students during placement, and language and cultural differences between international students and the workplace in the host country (30). |
|  | *Barrier to research dissemination (e.g., outreach and media coverage)* | Language was a barrier to research dissemination and networking during a collaborative experience in the UK, where culturally diverse participants interpreted specific concepts and ideas according to their own context (31). |
|  |  | Dissemination of information on agroforestry innovations was impeded due to language barriers in Sulawesi, Indonesia, as most farmers only speak local languages (32). |
|  |  | Language and cultural differences pose a barrier to the dissemination of indigenous knowledge in the form of storytelling (33). |
|  |  | Having English as the “International Language of Science” allows for the access of global scientific literature but creates a linguistic barrier for non-native English speakers, who are left out when disseminating their research (34). |
| **Policies** | *Inaccessibility to policy documents* | One third of the management plans identified for an assessment of strategic planning, zoning, impact monitoring, and tourism management at Natural World Heritage Sites were excluded from the analysis due to language barriers (19). |
|  | *Barrier to effective policy agreements among countries (e.g., bilateral agreements)* | Having a common official language between countries with established policy agreements had a statistically significant effect on reducing the tonnage of waste shipments (35) |
|  | *Language barriers between scientists and policy makers* | Language disparities can lead to poor communication between scientists and policy-makers (36). |
|  |  | Time available to read papers, difficulty in understanding technical language and reading in English have been recognized as a barrier to access scientific literature for Brazilian policy makers (37). |
| **Conservation activities** | *Barrier to developing collaborative actions* | Language barriers impede collaborative conservation activities between countries in the Mediterranean (11). |
|  |  | Differences in language represent a challenge for decision-making leading to agreements on multi-year resource allocation at National Parks in two transboundary regions in Europe, Italian-Slovenian and German-Czech borders (10). |
|  | *Extra costs (time/finance) for the multi-lingualisation of materials (websites, leaflets etc)* | Educational kits with information about the ecology and conservation of Critically Endangered spoon-billed sandpipers were translated to five different languages to improve the outreach of the conservation message (https://www.eaaflyway.net/spoon-billed-sandpiper-teaching-kit-available-for-free-download/). |
|  |  | Strategies to overcome language barriers in a collaborative decision analysis in two transboundary conservation regions in Europe included hiring multilingual staff (10). |
|  | *Inaccessibility to relevant information* | Differences in language in National Park planning documents represented a challenge for collaborative approaches for decision-makers in two transboundary conservation regions in Europe (10). |
|  |  | The dissemination of indigenous ecological knowledge and established conservation strategies in Australia is hindered by the fact that indigenous ranger groups, especially those in remote regions, who often do not speak English as their first language, are forced to adopt non-Indigenous forms of monitoring (38). |
| **General public** | *Difference in awareness due to cultural differences* | Local culture (a broad-scale consumption of the species) represents a major threat to yellow-breasted bunting, causing a population collapse of the species (39). |
|  |  | Use of indigenous ecological knowledge for contemporary land management has been limited by language barriers and cross-cultural awareness in Australia (38). |

Several studies have assessed the relationships between species diversity and linguistic diversity at local (*12*) to global scales (*13, 14*). However, despite growing evidence of the conservation consequences of language differences within species distributions, it remains unknown where such negative consequences of language barriers might be expected, and for which species. Here we investigate the number of languages spoken within the distribution of each of 10,863 extant bird species and discuss the ramifications of this for conservation. We focus on birds because (i) many bird species migrate, with their distribution spanning multiple countries, (ii) a wealth of ecological knowledge, especially detailed information on distribution is available (*15*), and (iii) a large number of transboundary conservation projects already exist (*16, 17*). We specifically aim to identify species with many languages within their distribution, and regions with high richness of such species, where language barriers could impede conservation.

## Results

On average, seven official languages are spoken within a species’ distribution, 16 for migratory species and three for threatened species. Additionally, 75.6% of the 10,863 extant bird species, 93.6% of the migratory species, and 55.5% of the threatened (59% of vulnerable (VU), 52.5% of endangered (EN) and 47.9% of critically endangered (CR)) species have two or more official languages within their distributions (Fig. 1).

**Figure 1.**
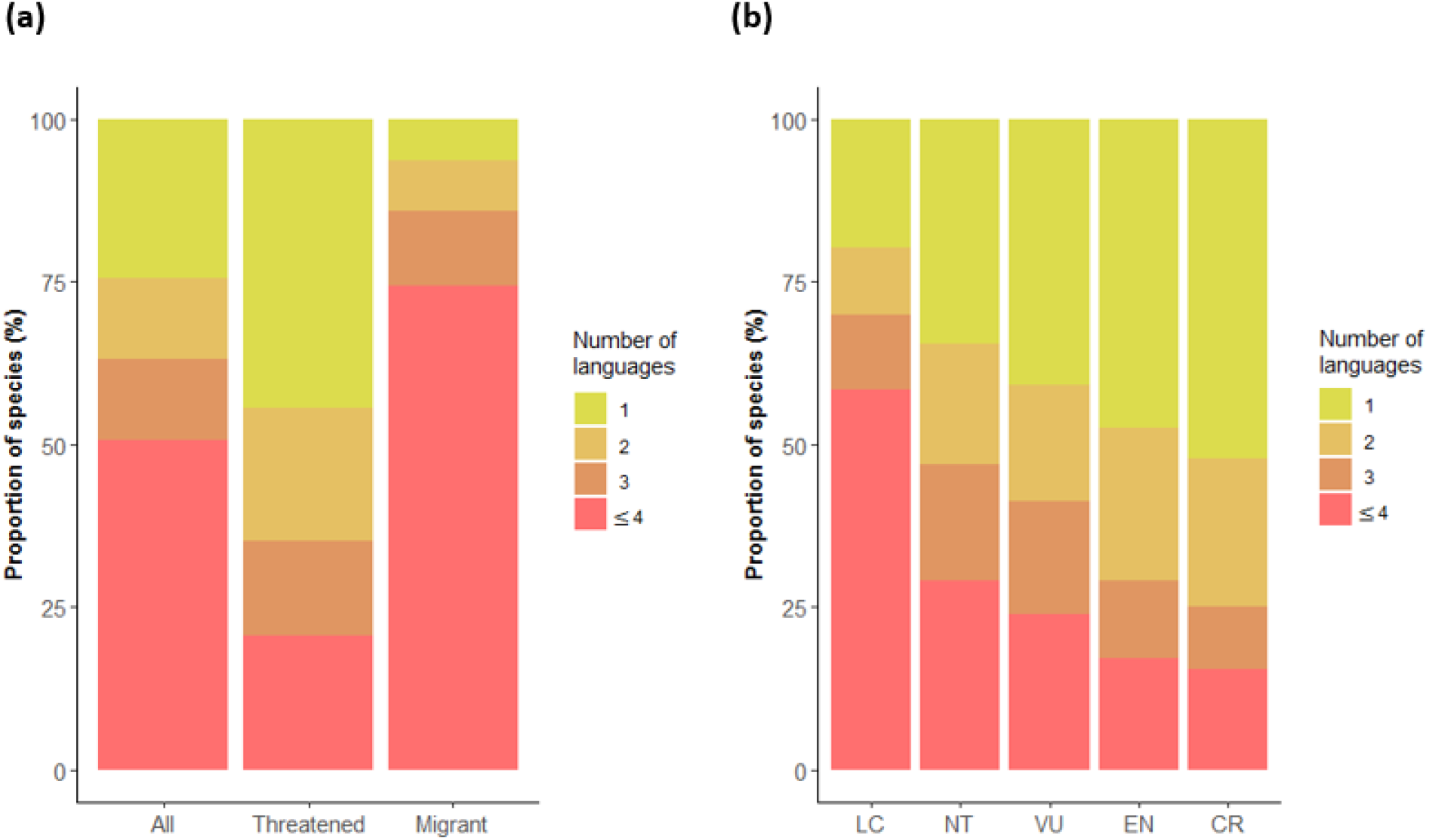
Language diversity among birds. **(a)** Number of official languages within the distributions of all bird species (n=10,863), threatened species (n=1427) and migratory species (n=1939). **(b)** Number of official languages spoken in the distributions of bird species by threat category (as assessed by the International Union for Conservation of Nature). See Figure S2 for data on the most spoken language in each country.

There is a strong positive relationship between the number of languages spoken within each species’ distribution and range size, and species with wide distributions have as many as 100 official languages spoken within their distribution (Fig. 2, Table S2). When controlling for the range size effect, threatened (CR and EN) and migratory species have significantly more languages spoken within their distributions, compared to non-threatened (LC) and non-migratory species (Fig. 2, Table S2). For example, Critically Endangered species with many languages within their distribution include Balearic shearwater (*Puffinus mauretanicus*, 25 languages), sociable lapwing (*Vanellus gregariu*s, 22 languages), and Rüppell’s vulture (*Gyps rueppelli*, 20 languages) (Fig. 2b). The results vary between taxonomic groups, with species in some orders, such as Strigiformes (owls) and Psittaciformes (including parrots, parakeets, lorikeets and macaws), having comparatively few official languages within their distribution (seven and three on average, respectively), with others, such as Ciconiiformes (including storks, herons, bitterns, ibises and spoonbills) and Charadriiformes (including waders, gulls and auks), having especially many languages (19 and 17 on average, respectively; Fig. 2c). The results were qualitatively the same based on the most spoken languages in each country (Fig. S3, Table S2).

**Figure 2.**
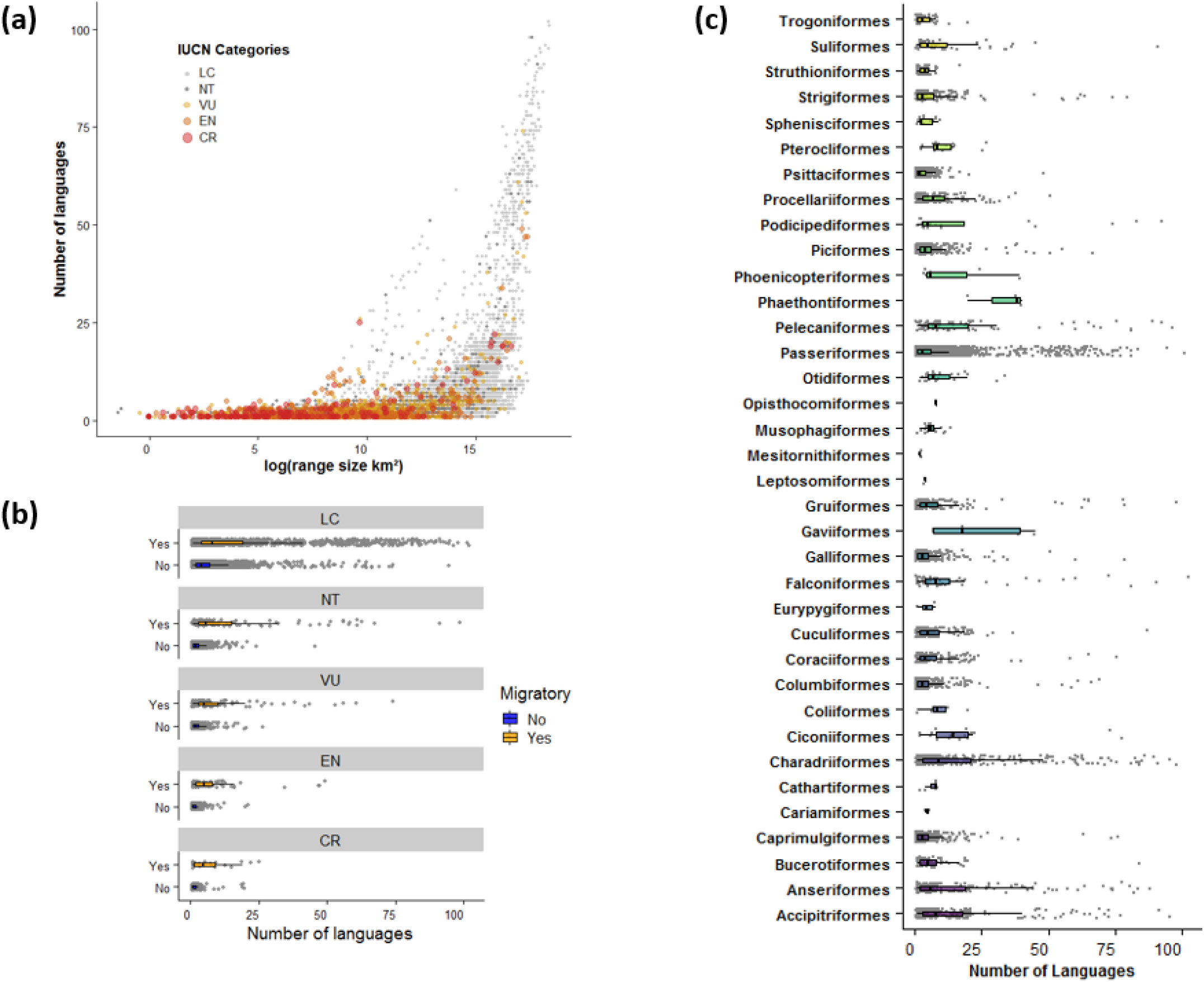
**(a)** Relationship between bird species’ distribution range size and the number of official languages within their distribution. International Union for Conservation of Nature (IUCN) threat categories are shown in different colours. Number of official languages spoken within each species’ distribution by **(b)** migratory status and IUCN threat categories, and by **(c)** taxonomic order. See Figure S3 for the same figure but based on the most spoken language in each country.

English, Spanish, French and Portuguese are the four languages associated with the most species; this pattern was consistent for all species, threatened species, and migratory species (Table S3, Fig. S4). Across all bird species, 45% have some area of their distribution associated with Spanish, 38% with English, 27% with Portuguese and 22% with French. For migratory species 67% were associated with English, 61% with Spanish, 42% with French and 38% with Portuguese. Finally, for threatened species, 23% were associated with Spanish, 16% with English, 16% with Portuguese and 12% with French (Table S3). However, 899 species associated with Spanish are not associated with any other languages and thus, when only species associated with two or more languages were assessed English was the language associated with the most species, for all species and for threatened species (Table S3). Geographically there is variation in the distribution of the species associated to the top six languages; In south America many species are associated with English, Spanish and Portuguese, in Africa with English, Kiswahili, Portuguese and French, and in Southeast Asia with Mandarin (Fig. 3; see https://translatesciences.shinyapps.io/bird_language_diversity/ for other languages’ results).

**Figure 3.**
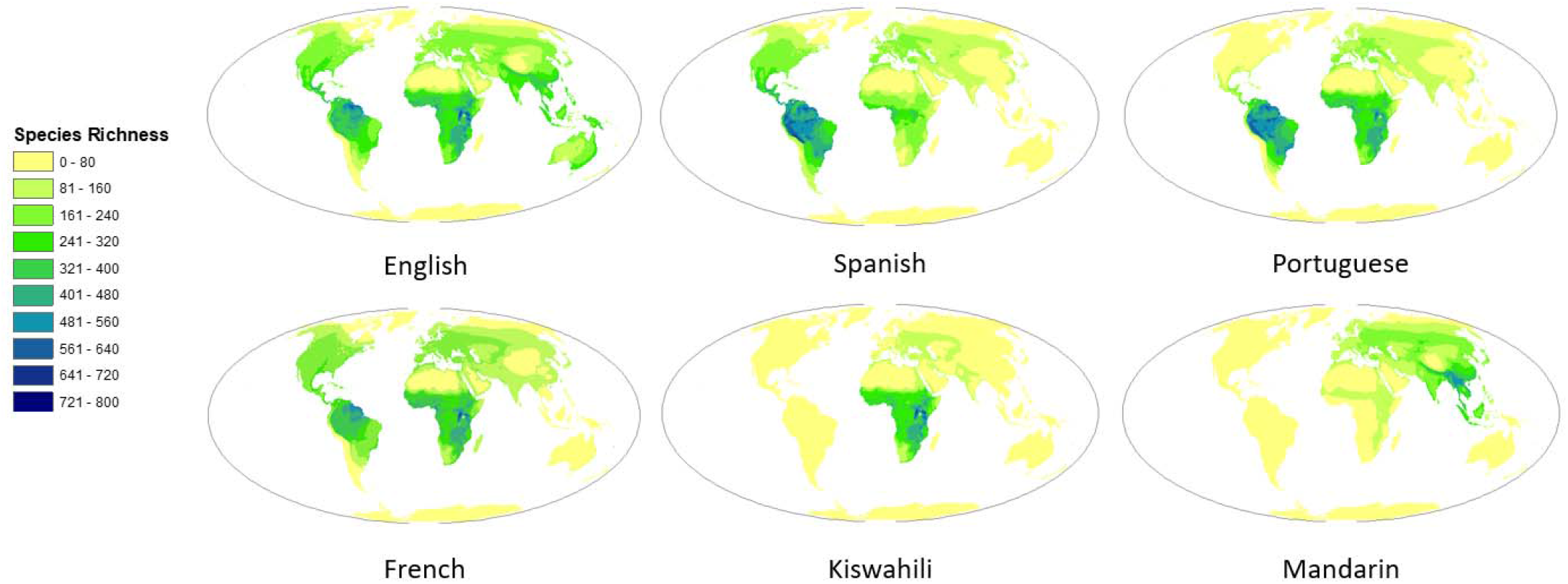
Species richness of birds associated with each of the top six official languages with the highest number of species. See https://translatesciences.shinyapps.io/bird_language_diversity/ for other languages’ results.

Especially many species with high numbers of languages spoken within species distributions were found in central and southern Africa, India, southeast China, eastern Europe, and Russia (Fig. 4a). A large number of threatened species with high language richness were found in Western and Central Asia as well as southern Russia (Fig. 4b). A similar pattern was found for migratory species with eastern Europe also being a hotspot of species with high language richness (Fig. 4c). The results remained qualitatively the same when using the most spoken language, instead of official languages, in each country (Fig. S2, 3, 4 & 5).

**Figure 4.**
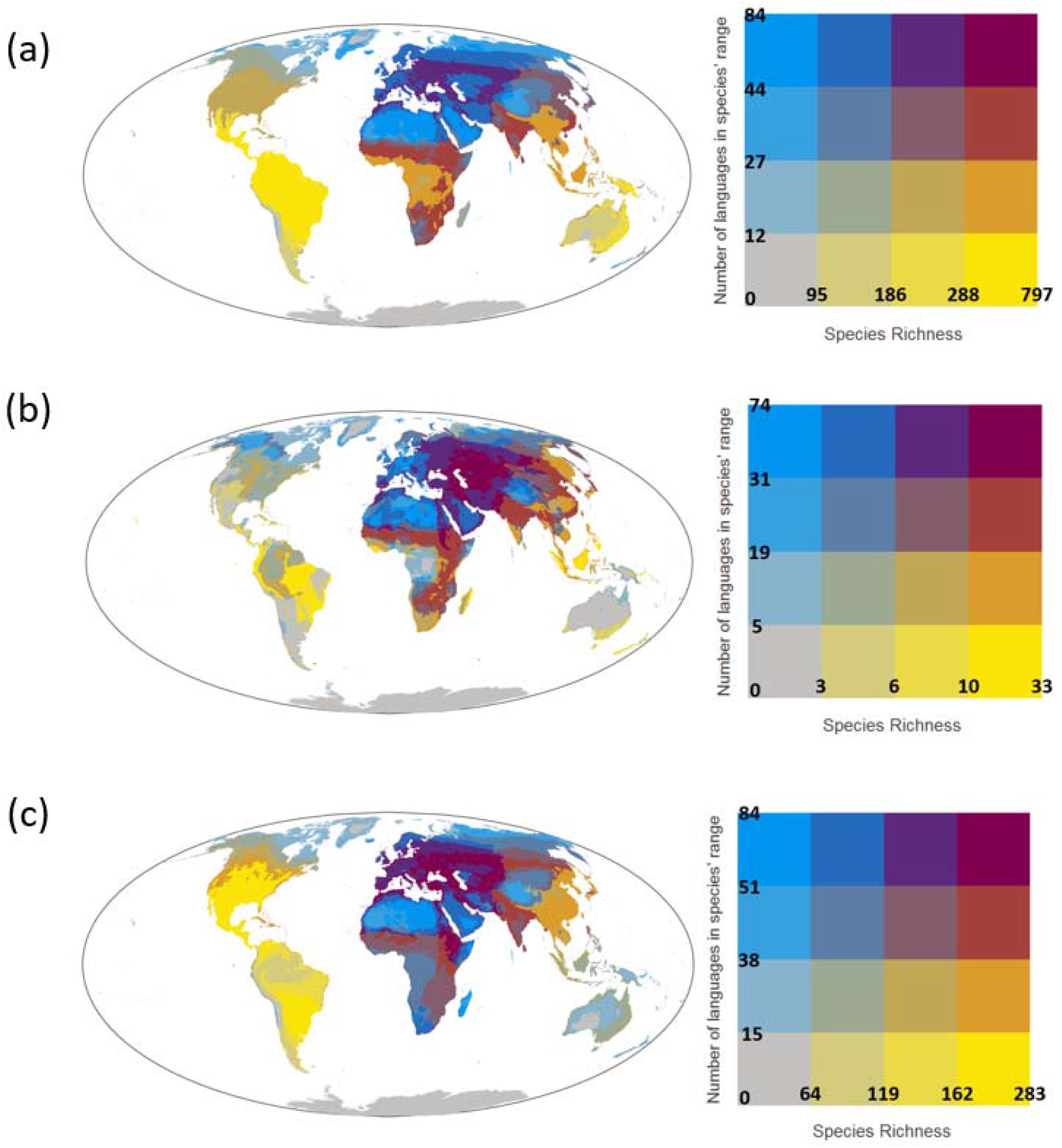
Bivariate maps showing the number of species (species richness) and the mean number of languages within the distribution of species found within each 30km × 30km grid cell for **(a)** all bird species, **(b)** threatened bird species, and **(c)** migratory bird species. The number of languages within each species’ distribution was calculated using the official languages in each country. See Figure S5 for the same figure but using the dataset of the most spoken language in each country.

## Discussion

Language differences are increasingly recognized as a barrier to transboundary conservation, and several international projects have been developing guidelines on how to overcome this barrier (*18, 19*). As summarized in Table 1, language differences can have serious consequences for conservation by, for example, posing barriers to the generation and transfer of scientific knowledge as well as the development of effective conservation activities and policies. Such negative consequences of language barriers are expected to be particularly severe in the conservation of species with multiple languages being spoken within their distribution. Our research provides important insights into where in the world and for which species conservationists are especially required to make extra efforts to overcome language barriers to improve bird conservation.

Our results reveal that threatened (CR and EN) and migratory species have more languages spoken within their distribution, when controlling for range size (Table S2). Additionally, Our results show that 217 bird species have 50 languages or more spoken within their distributions (Table S4) and that more than two thirds of all species, half of the threatened and almost all migratory species are associated with two or more languages (Fig. 1). These results, together with the multiple ways that language barriers can affect conservation (Table 1), highlight the potentially serious consequences of language barriers in bird conservation, especially for migratory and threatened bird species. For example, the distribution of the common pochard (*Aythya ferina*), which is classified as Vulnerable by the IUCN, spans 108 countries in Europe, Russia, Asia, and north Africa, where a total of 75 official languages are spoken. This means that scientific information on this species (including peer-reviewed papers and grey literature) can be scattered across those different languages, and successful conservation of the species may depend on effective collaboration and policy agreements among people with diverse linguistic and cultural backgrounds. Species in the orders Ciconiiformes and Charadriiformes have an especially high number of languages spoken within their distributions. For example, the Critically Endangered spoon-billed sandpiper (*Calidris pygmaea*) has nine different languages spoken within its distribution. For this species, educational kits with information about the species ecology and its conservation have already been translated to five different languages to improve the outreach of the conservation message (https://www.eaaflyway.net/spoon-billed-sandpiper-teaching-kit-available-for-free-download/), demonstrating the work required to address language barriers in conservation. The conservation of species associated with many languages will likely require such coordinated efforts among stakeholders with different cultural and linguistic backgrounds, for example through incorporating action plans to overcome language barriers in relevant policy agreements, such as those in the Convention on Migratory Species (*20*).

Even though one third of the bird species globally have English spoken within a part of their distribution, other languages are also associated with a large number of species in certain regions, such as Spanish and Portuguese in South America, Kiswahili in Africa, and Mandarin in South East Asia. These languages could be key to conservation research, policies, and practices in those regions. For example, important information related to species ecology and conservation is often available in non-English languages (*6*), which is however usually omitted when conducting conservation research and generating conservation plans. The omission of such non-English-language information can bias inferences of ecological analysis (*21*), which in turn can cause suboptimal conservation decisions. Effective conservation of bird species would require synthesizing scientific information and transferring generated knowledge in these key languages, and our results provide practical information on which species would benefit from multilingual assessments and which languages are key to those species (see Table S4 and Fig. 4, see https://translatesciences.shinyapps.io/bird_language_diversity/ for other languages’ results).

Overcoming language barriers will play an important role in areas with a large number of species with many languages spoken within their distribution. These regions include central and southern Africa, India and Southeast Asia as well as Kazakhstan, southern Russia and Western Asia for threatened and migratory species. Challenges for bird conservation in these regions include a need to reconcile perspectives and interests among extremely diverse stakeholders, as species in these regions have, on average, up to 84 different languages spoken within their distributions. Establishing cross-national associations, such as the European Bird Census Council (https://www.ebcc.info/), in these regions would be an effective approach for coordinating monitoring and conservation efforts and achieving consensus decisions among countries. Other areas where species associated with particularly many languages are found are Europe, north Africa, western Asia and north Russia; again up to 84 languages, on average, are spoken within the distribution of species found in those regions. Although these regions did not show the highest richness of such species, this does not diminish the importance of proactively accounting for language barriers in conservation initiatives in these areas. For example, the United Nations Barcelona Convention has developed guidelines on conservation of Mediterranean seabirds that promote coordinated actions between countries with different language backgrounds. The Mediterranean Small Island Initiative also aims to facilitate collaborations between ten different countries in the region (*17*). Such initiatives would benefit from the creation of guidelines to overcome language barriers between the parties involved (*11*).

While our analysis is, to our knowledge, the most comprehensive assessment of the identity and number of languages spoken through the distribution of bird species globally, the way this association was measured has some caveats. The presence of a species on a particular country does not imply that all of the official languages spoken in that country are spoken within that species distribution. Additionally, the fact that multiple languages are spoken through the distribution of a species does not imply that all the scientific information is being generated in different languages through the species distribution. Education systems in many countries promote learning of multiple languages (including English) that are different from the official or most spoken one in the country (*22*) (*23*). Future research is needed to understand the ability of people to work across language barriers and how it varies geographically, and also to identify particular species with low compatibility between the languages spoken within their distribution, areas with an especially large number of such species, and languages that generate such incompatibility.

The global community has a joint responsibility to address the biodiversity crisis and avoid further species extinctions, which requires an effective transfer of knowledge and information between countries with diverse linguistic and cultural backgrounds. In Table 1 we identified four different pathways through which language barriers affect conservation: 1) scientific research, 2) policies, 3) conservation activities and 4) general public. Here we provide potential solutions to overcome such ramifications of language barriers in conservation. A way to improve the transfer of scientific research and to overcome language barriers when generating and executing conservation policies is to promote the multilingual transfer of relevant information, ideally through a clear, concise and easy-to-use translation protocol (*24*), especially for species with many languages spoken within their distributions and in areas where those species are found. This can be done by, for example, providing translations of relevant scientific papers and policy documents in multiple, relevant languages. Using information sourced from multiple languages, especially languages associated with the species being assessed, and actively engaging with scientist and politicians with different language and cultural background would also increase the access to otherwise omitted information. This improves the quantity and quality of the knowledge on the ecology and conservation of the species, which in turn facilitates the generation and execution of more effective conservation policies. On the other hand, stimulating multilingual conservation activities, such as the ones implemented in the program “Birds without borders” (https://www.birdlife.org/africa/projects/conservation-migratory-birds-cmb), as well as promoting the translation of critical conservation information on target species into clear and brief documents for the general public would improve the success of conservation actions and the outreach of information on how to avoid the extinction of those species. Our analysis has shown species and areas with significant challenges of language barriers to conservation and we have provided some potential solutions for these challenges. To implement these solutions and overcome these barriers there is a need for political will, local support and sufficient resourcing (*4, 6*).

## Acknowledgments

We are grateful to H. P. Possingham, D. Biggs, L. Sonter, and the Fuller lab group for providing constructive feedback and discussions around elements of this study.

## Author contributions

T.A. and P.J.N. designed the research. P.J.N. performed the analysis with help from S.C.A and T.A. the Shiny app was developed by B.K.W. The manuscript was written by P.J.N with help from S.C.A., B.K.W., M.C., J.R.A., R.A.F., and J.E.W.

## Data Availability

Supplementary Table 1,3 & 4 will be made available upon request, previous to its deposition in an open-access repository with the peer-reviewed version of this study. Requests should be sent to the corresponding author.

## Supplementary information

### Materials and Methods

#### Bird species data

We obtained species distribution maps for the birds of the world from Birdlife International and Nature Serve (*1*). We considered parts of each species distribution coded as “extant” for presence and “native” and “reintroduced” for origin. In the case of migratory species, all seasonal sections of the distribution were considered. Additionally, we obtained information on taxonomic classification, threat status and type of migratory characteristics for each species from Birdlife (*1*). Species were divided by conservation status [ie., threatened (VU, EN and CR) and not threatened (LC and NT)] and migratory status [(i.e., Full migrants or not (the latter comprising non-migratory, altitudinal migrants and nomadic species)], and results were aggregated for these groups and for each bird species. The area of each species range distribution was calculated (in km²) Using PostGIS version 3.0.2 (*2*).

#### Data on languages of the world

We compiled information on the official and most spoken languages of each country of the world. Official languages are the ones used by a country or jurisdiction for governmental and legal purposes while the most spoken language is the one that the largest proportion of the population of a country or jurisdiction speak. We used the World Fact Book from the United States Central Intelligence Agency (*3*) as a primary source, but additional sources were used as needed (see Table S1). For the official languages we listed all the languages that each country states as official. Spain (five official languages), Ethiopia (five) and South Africa (11) were the only countries with more than four official languages so for those countries the top four official languages with the highest number of speakers were used. For disputed regions with official information available, such as Kashmir, the most commonly spoken languages in the region were used. This information was gathered from additional sources (Table S1). For Antarctica no official language was assigned. We also used the World Fact Book to identify the most spoken language in each country or jurisdiction. If this information was not available, the language recorded as “lingua franca” in the World Fact Book was selected. For Antarctica and Kashmir no language was assigned as most spoken.

#### Calculation of number of languages in species distribution and bird species richness

First, we determined the identity of the countries each species distribution overlaps with and the official and most spoken languages of those countries. Those languages were assigned to each species. Then we estimated bird species richness using a global 30 km × 30 km grid. This has been identified as an optimal resolution for reducing the effects of commission errors (where species are thought to be present but are not) when working with global species distribution maps (*4*). Grid cells that straddle more than one country were split through the country borders into sub-units for each country. The number (i.e., species richness) and identity of the species present in each grid cell was determined.

#### Mapping areas of high numbers of bird species with many languages within their distribution

Finally, by using the identity of the species present in each grid cell and the information on the number of languages spoken in the distribution of each species, we calculated the mean number of languages spoken in the distributions of the species present in each grid cell (Fig. S1). Using this information, we were able to identify areas in the world with high numbers of bird species with many languages within their distribution. Spatial data were analyzed in a Mollweide equal area projection in ESRI ArcGIS version 10.4 (*5*) and PostGIS version 3.0.2 (*2*), and statistics were calculated in R statistical language version 3.5.1 (*6*).

#### Statistical analysis

To investigate factors explaining the number of languages spoken within each species’ distribution, we performed generalized linear mixed models (GLMMs) assuming a negative binomial distribution with the number of (either official or most spoken) languages spoken within each species’ distribution as the response variable, log_10_-transformed distribution range size (km²), migratory status (non-migrant as the reference category), and conservation status (Least Concern (LC) as the reference category) as the explanatory variables, and the order of each species as a random factor. The GLMMs were implemented using the package lme4 in R (*7*).

**Table S1,3 & 4 are in excel format**

**Table S1.** List of official and most spoken languages for each country in the world.

**Table S3.** Number of bird species (n= 10863) associated with each of the official and most spoken languages of each country in the world.

**Table S4.** Number of official and most spoken languages associated with each bird species assessed (n=10863).

**Table S2.**
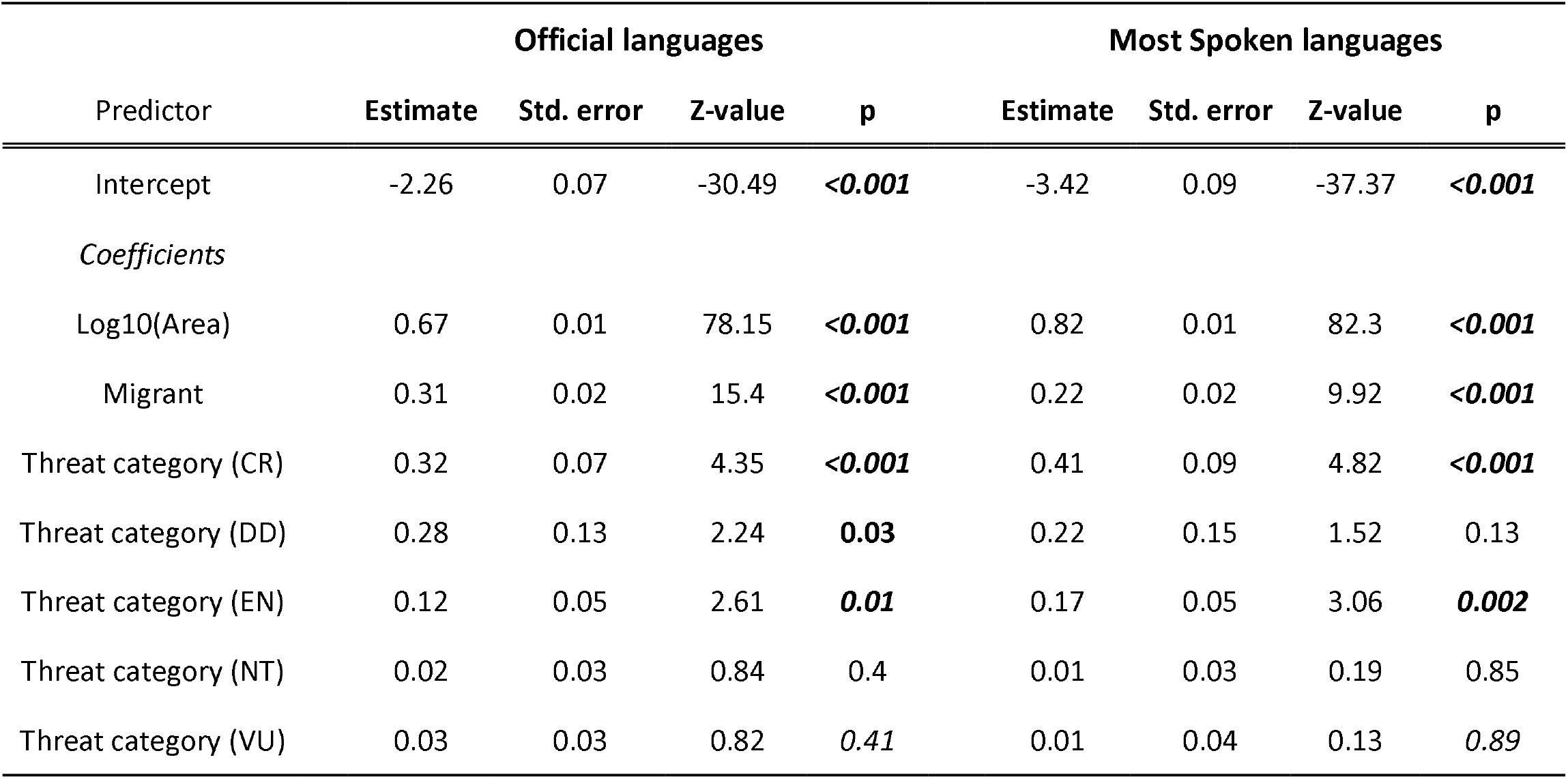
Results of negative binomial generalized linear mixed models (GLMM) to explain the number of (official or most spoken) languages spoken within bird species distribution (the response variable) using the three explanatory variables: log_10_-transformed distribution range size (km^2^), migratory status (non-migrant as the reference category), and IUCN threat categories (Least Concern as the reference category). The order of each species was also incorporated in the models as a random factor. Statistically significant p-values (p < 0.05) are indicated in bold.

**Figure S1.**
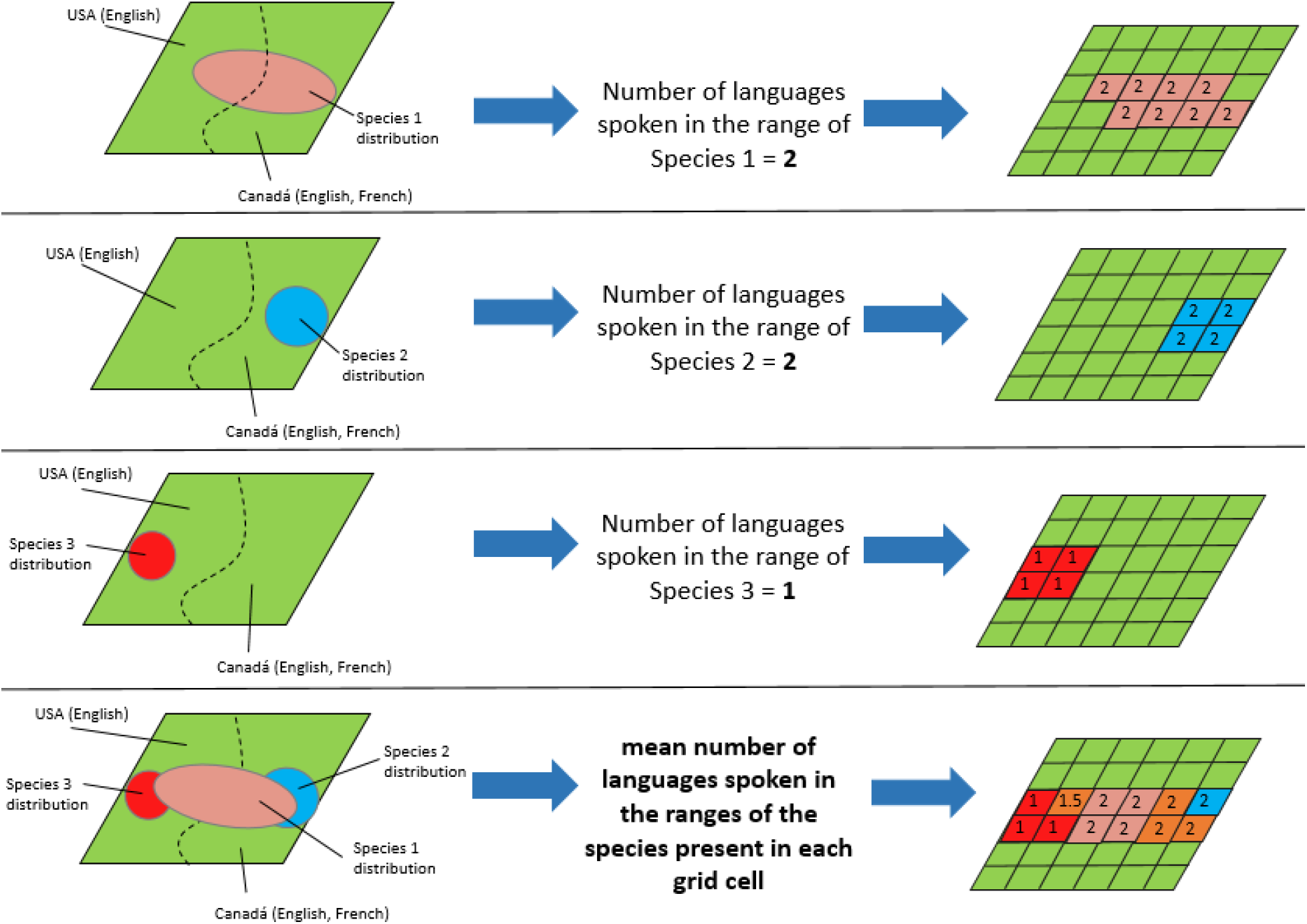
Methodological framework for mapping mean linguistic diversity across all species within each grid cell

**Figure S2.**
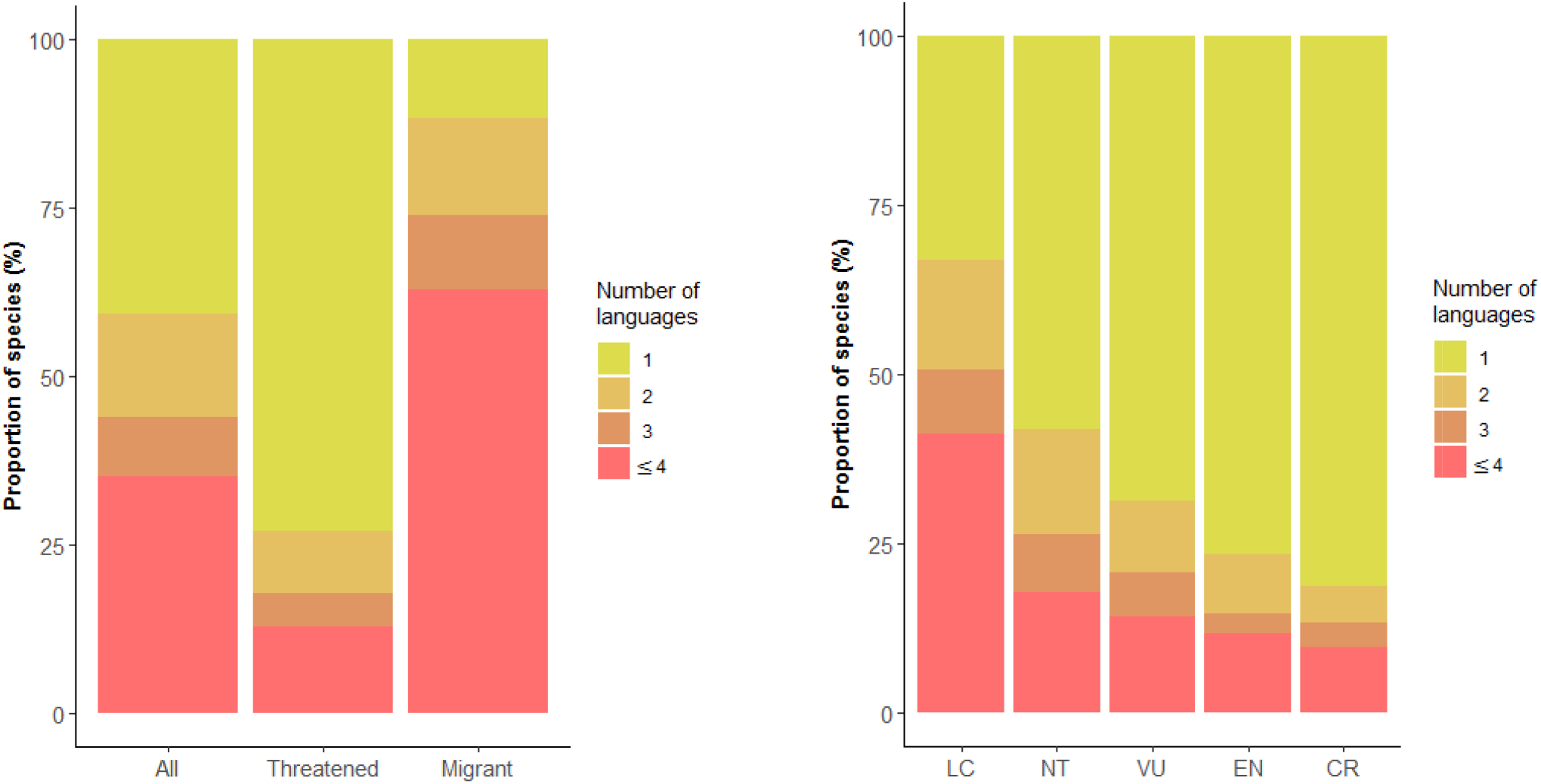
Language diversity among birds. **(a)** Number of most spoken languages within t distributions of all bird species (n=10,863), threatened species (n=1427) and migratory species (n=1939). **(b)** Number of most spoken languages spoken in the distributions of bird species by threat category (a assessed by the International Union for Conservation of Nature).

**Figure S3.**
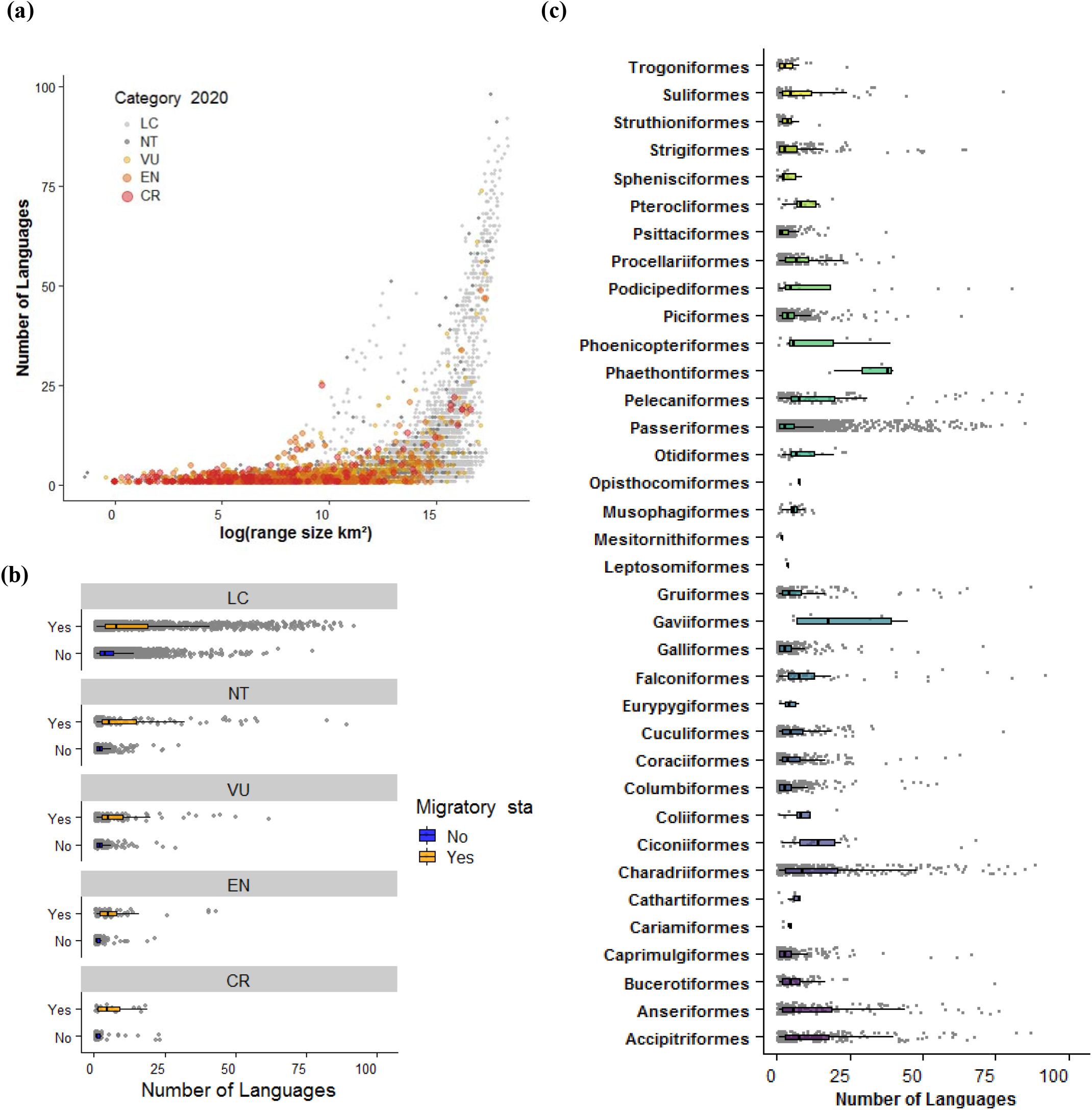
**(a)** Relationship between bird species’ distribution range size and the number of language within their distribution. International Union for Conservation of Nature (IUCN) threat categories are shown in different colours. Number of languages spoken within each species’ distribution by **(b)** migratory status and IUCN threat categories, and by **(c)** taxonomic order. This analysis was done using the dataset of most spoken language in each country.

**Figure S4.**
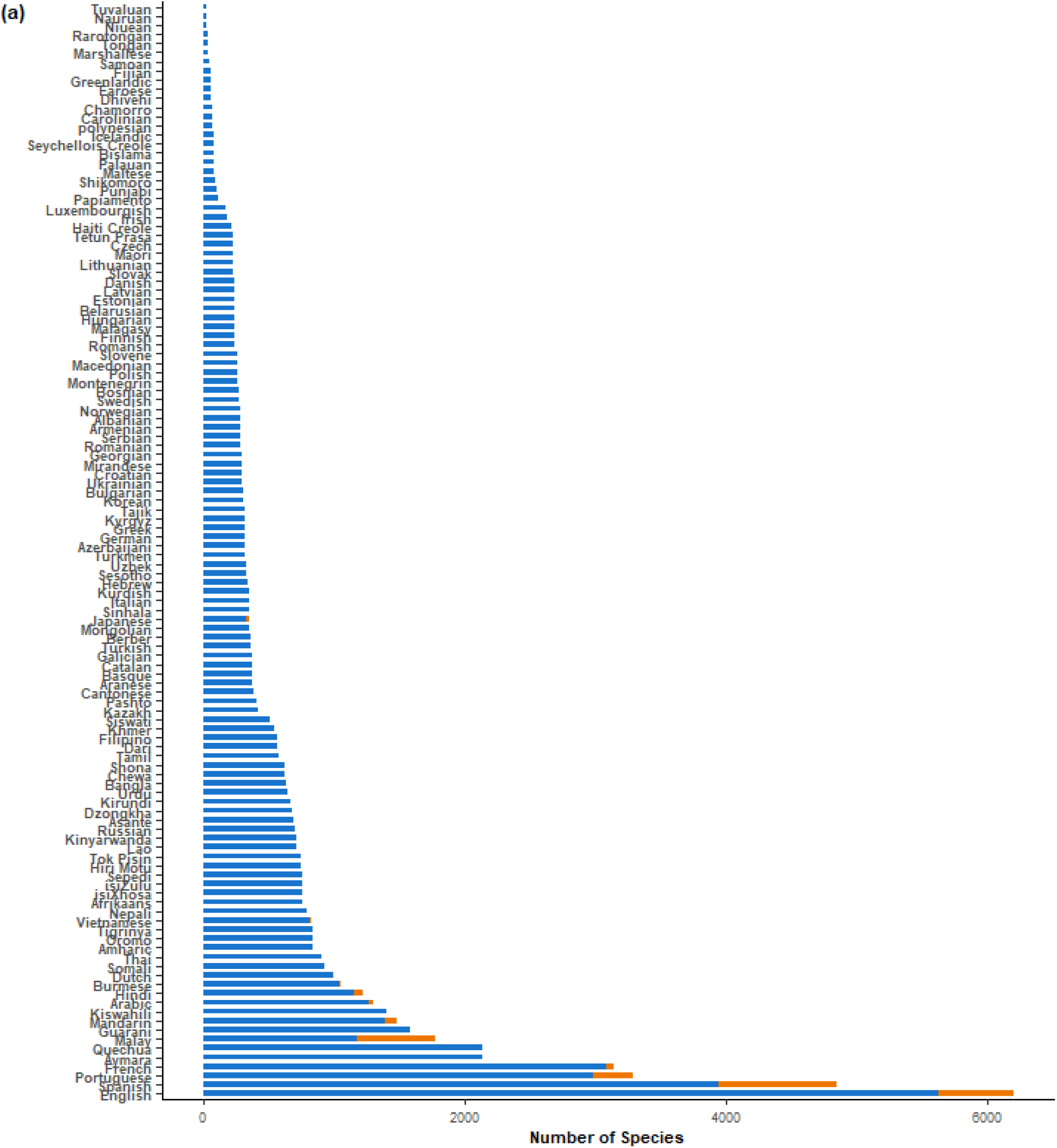
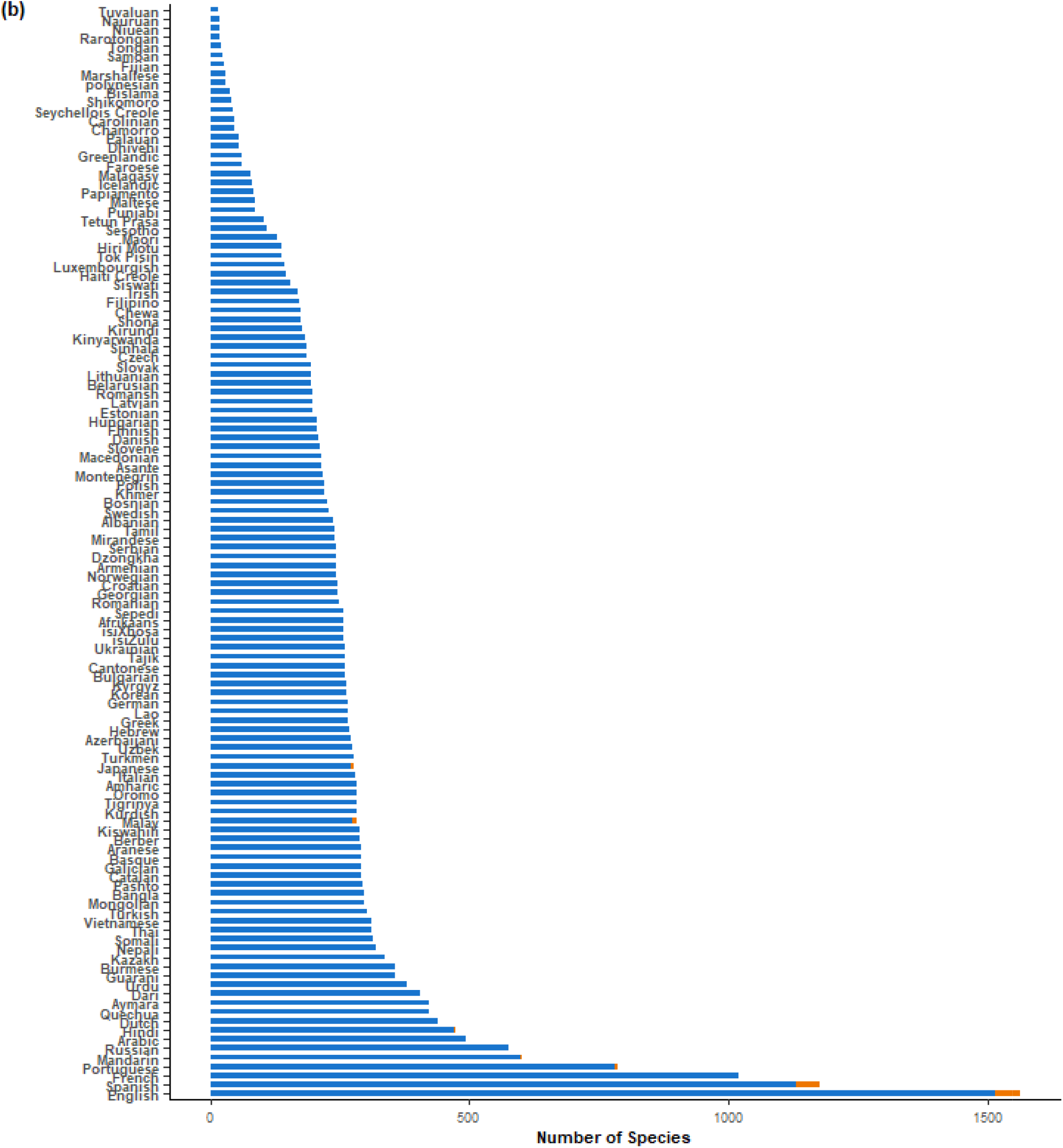
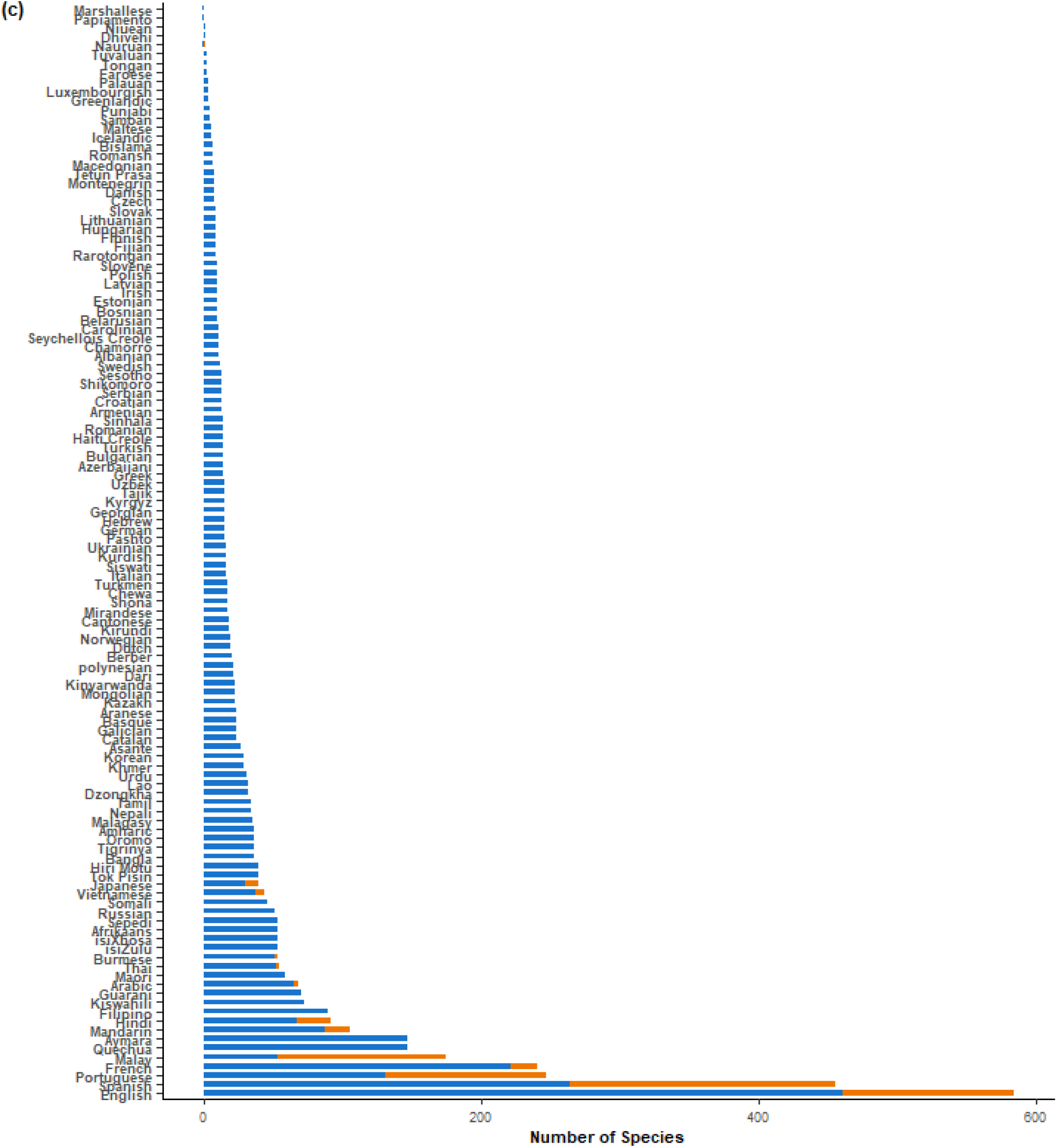
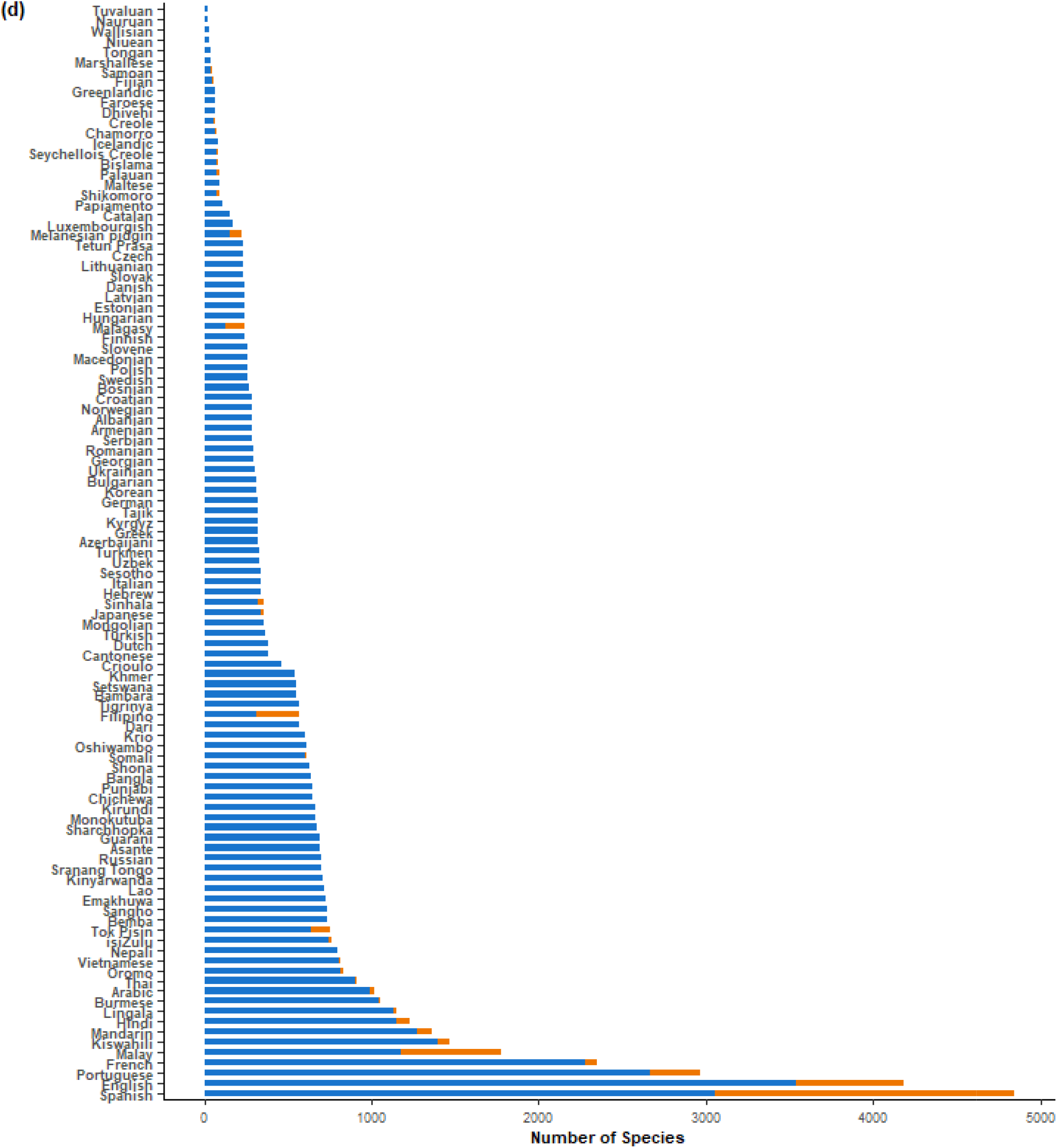
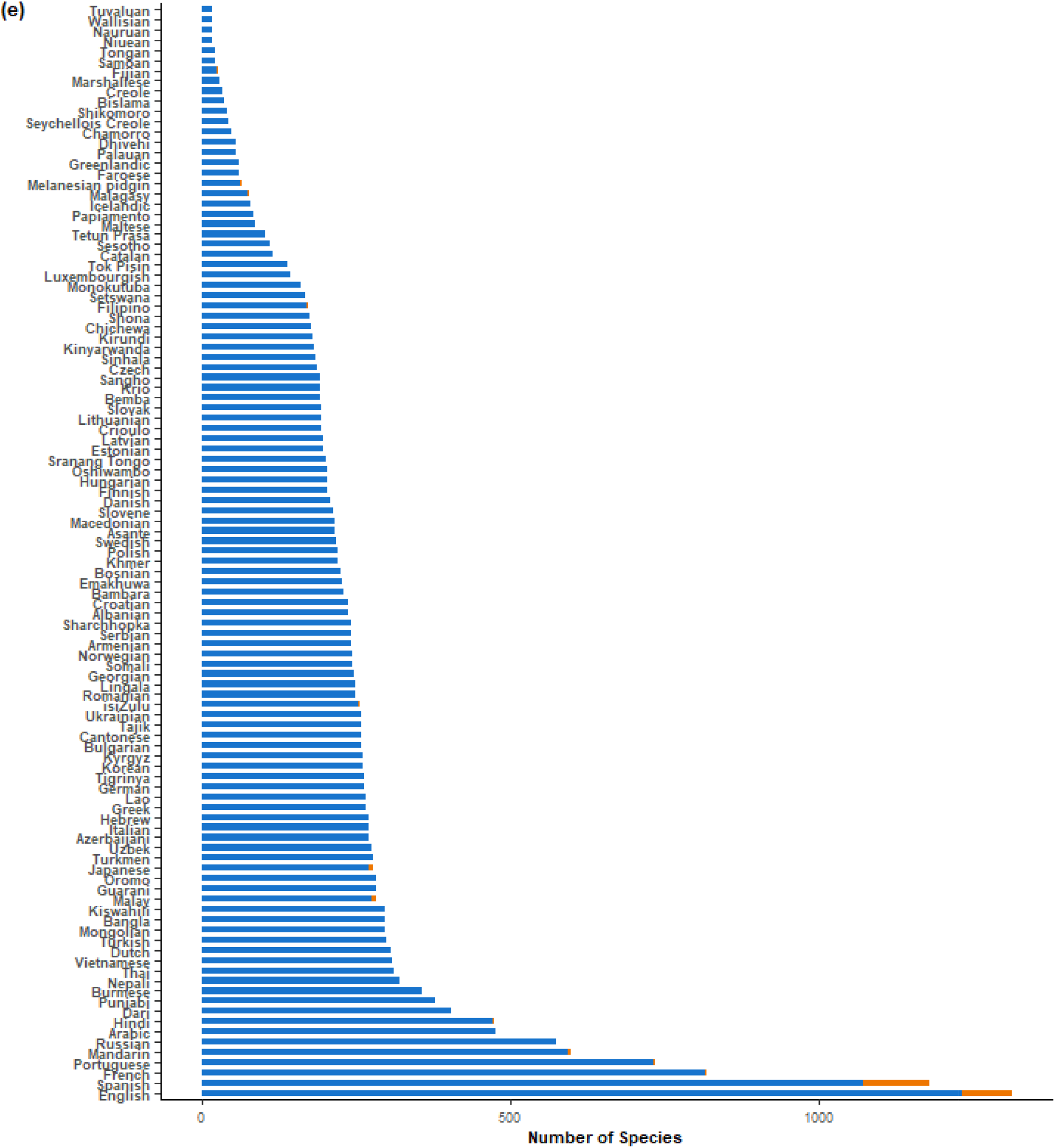
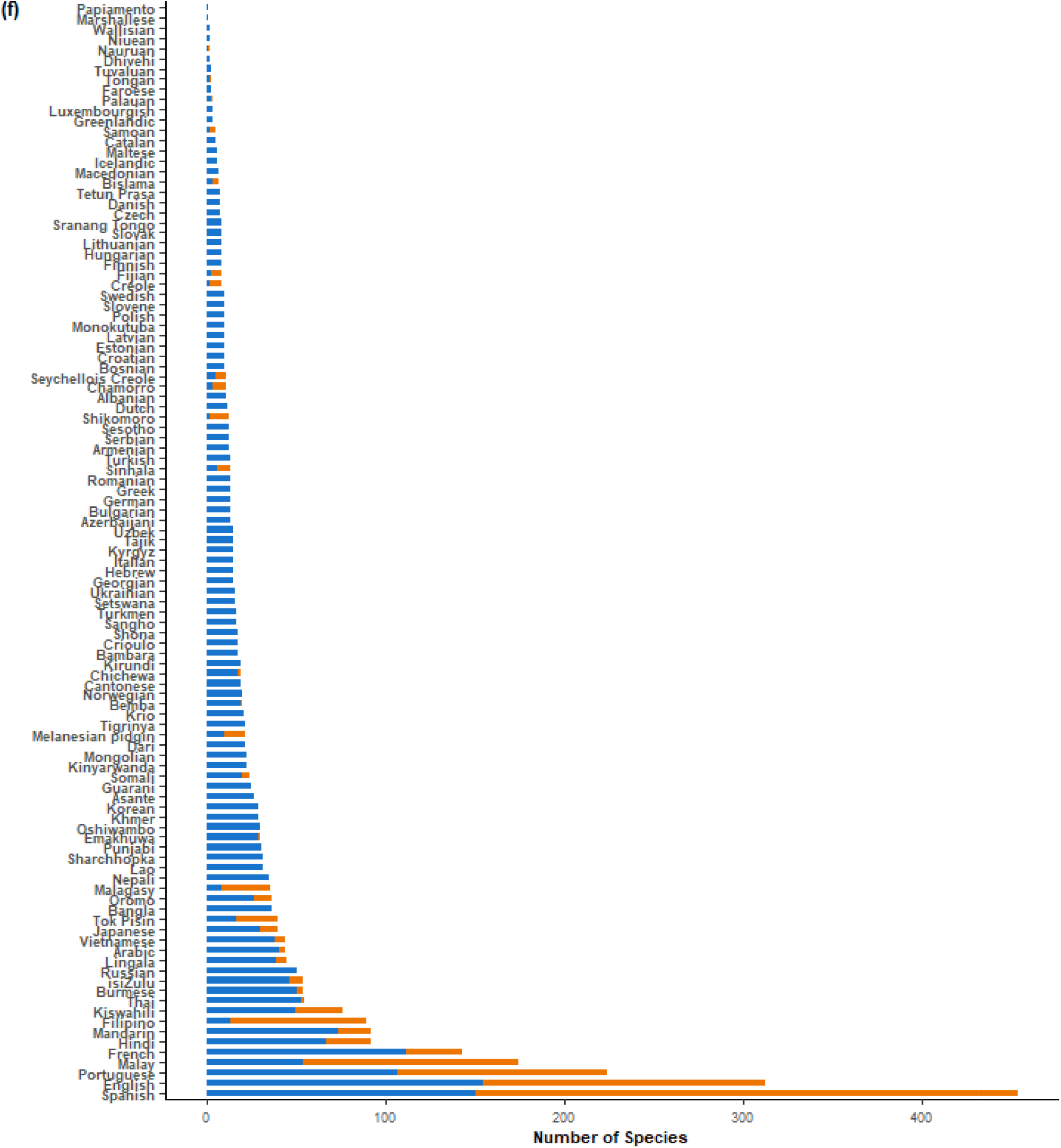
Number of bird species associated with a particular language. Number of official language associated with **(a)** all species (n=10,863), **(b)** migratory species (n=1,939), and **(c)** threatened specie (n=1,427). The same analysis but for most spoken languages in each country for **(d)** all species, **(e)** migratory species, and **(f)** threatened species. The number of species associated only with the language i shown in orange and the number of species associated with the language and one or more other language is shown in blue.

**Figure S5.**
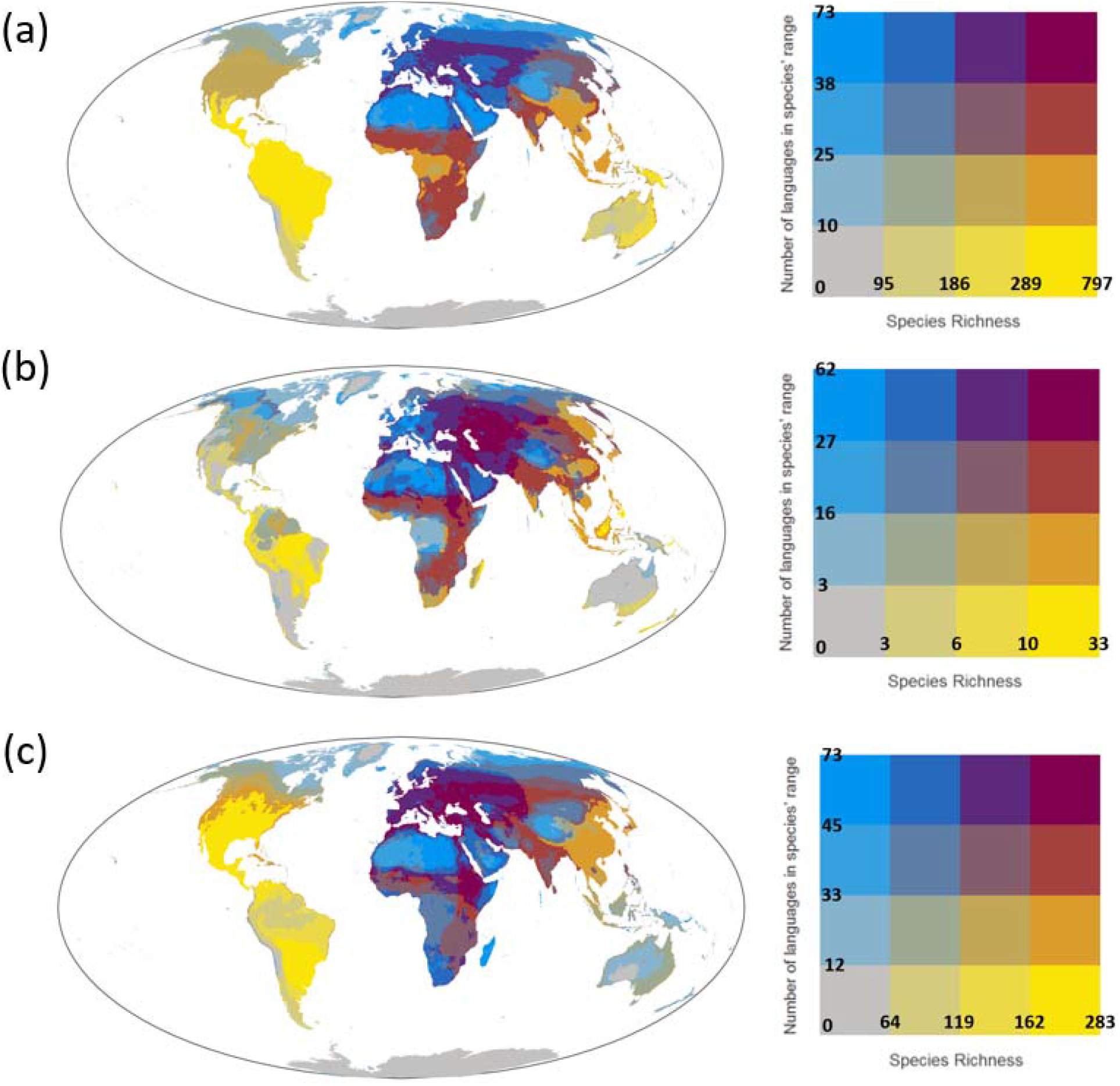
Bivariate maps showing the number of species (species richness) and the mean number of languages within the distribution of species found within each 30km × 30km grid cell for **(a)** all bird species, **(b)** threatened bird species, and **(c)** migratory bird species. The number of languages within each species’ distribution was calculated using the most spoken languages in each country.

## Notes

### Competing Interest Statement

The authors have declared no competing interest.

